# Periplasmic stress contributes to a tradeoff between protein secretion and cell growth in E. Coli Nissile

**DOI:** 10.1101/2023.01.09.523330

**Authors:** S Emani, A Kan, T Storms, S Bonanno, J Law, S Ray, N Joshi

## Abstract

Maximizing protein secretion is an important target in the design of engineered living systems. In this paper, we characterize a tradeoff between cell growth and per cell protein secretion in the curli biofilm secretion system of E Coli Nissile 1917. Initial characterization using 24-hour continuous growth and protein production monitoring confirms decreased growth rates at high induction leading to a local maximum in total protein production at intermediate induction. Propidium iodide staining at the endpoint indicates that cellular death is a dominant cause of growth reduction. Assaying variants with combinatorial constructs of inner and outer membrane secretion tags, we find that diminished growth at high production is specific to secretory variants associated with accumulation of protein containing the outer membrane transport tag in the periplasmic space. RNA sequencing experiments indicate upregulation of known periplasmic stress response genes in the highly secreting variant, further implicating periplasmic stress in the growth-secretion tradeoff. Overall, these results motivate additional strategies for optimizing total protein production and longevity of secretory engineered living systems.

## Introduction

Over the past few decades, engineered living systems have emerged as a revolutionary technology for accomplishing complex tasks such as chemical synthesis, autonomous detection, and therapeutic drug delivery [1]. Protein secretion is an important goal for engineered living systems, as it enables organisms to respond to and influence their extracellular environment, which is relevant for self-assembly of living materials, sensory circuits and interorganismal communication [2-4]. For many of these tasks, bacteria which produce high protein yield with high efficiency are desirable. Thus, improving the secretion efficiency and productivity of bacterial systems is an important goal for engineered living systems research.

Any biological system engineered to perform a specific task faces an inherent trade-off between productivity and survival. One can only expect to increase activity in a single cell up to a point before metabolic constraints and cell stress limit the cell’s ability to reproduce. When protein secretion is involved, the burden of increased production is exacerbated by the energetic demands of cross-membrane transport and interference with other vital secretory pathways [5]. Understanding the relationships between cellular growth and per-cell protein secretion is therefore crucial to optimizing the total secretory capacity of an engineered system. Revealing the mechanisms of growth impairment caused by increased secretion could enable the design of bacterial strains that are more resistant to such stress or feedback systems that enable self-optimization of per-cell protein production [6]. Moreover, cell survival is often an independently important target for engineering, since it is critical for stable long-living systems (e.g. colonization of a host organism for probiotics) [7].

Protein secretion in *E. coli*, like in other gram-negative bacteria, can proceed through several known pathways [8]. Although the double membrane structure complicates these pathways compared to gram-positives, *E. coli’s* high engineerability and therapeutic potential as a natural probiotic make it a valuable target for secretion optimization [9, 10]. We chose to focus on optimizing the Type VIII secretion system because of its association with biofilm formation, during which large quantities of extracellular protein fibers called “curli” are produced [11]. Curli fibers are composed primarily of a single protein monomer, CsgA, which self-assembles into filaments anchored to the outer membrane. CsgA utilizes the well-characterized Sec inner membrane transport channel to traverse the inner membrane, and a dedicated protein channel encoded by the curli operon (composed of csgE and csgG) for outer membrane secretion [12]. By repurposing the secretory apparatus used by curli and substituting csgA with a fluorescent reporter protein, we built a model system for optimizing protein secretion and investigating growth-secretion tradeoffs in *E. coli*.

In this paper, we investigate the relationship between growth and protein secretion in *E. coli* Nissile 1917 by secreting super-folder GFP (sfGFP) through the native curli secretion machinery. sfGFP was chosen as the primary protein of interest due to its globular protein structure and its easy quantifiability with a fluorescence assay. First, we empirically characterize the tradeoffs between growth rate and per-cell protein production. We then utilize a series of alterations to the secretion apparatus to identify the mechanism of growth depression and sequence the RNA transcripts associated with secretion stress. Our work illuminates mechanistic aspects of secretion stress and as such identifies valuable targets for secretion optimization.

## Results

The tradeoff between cell growth and protein output was investigated in a modified strain of *E. coli* Nissle 1917, PBP8, which has the curli operon replaced with a chloramphenicol resistance gene (∆csgBACDEFG::Cm^R^). The strain was transformed with a plasmid vector encoding the curli machinery with sfGFP substituted for the CsgA monomer. As with CsgA, sfGFP was tagged with N-terminal signal sequences for inner and outer membrane secretion, called the Sec and N22 tags, respectively [13]. The sfGFP gene was placed under the control of an IPTG inducible promoter, while the channel proteins (csgE, csgF, and csgG) were constitutively expressed as the native polycistronic operon (Figure 1a). We also constructed variant plasmids that removed one or both of the N-terminus secretion tags on the sfGFP, with or without the csgEFG operon (Figure 1A).

**Figure 1.**
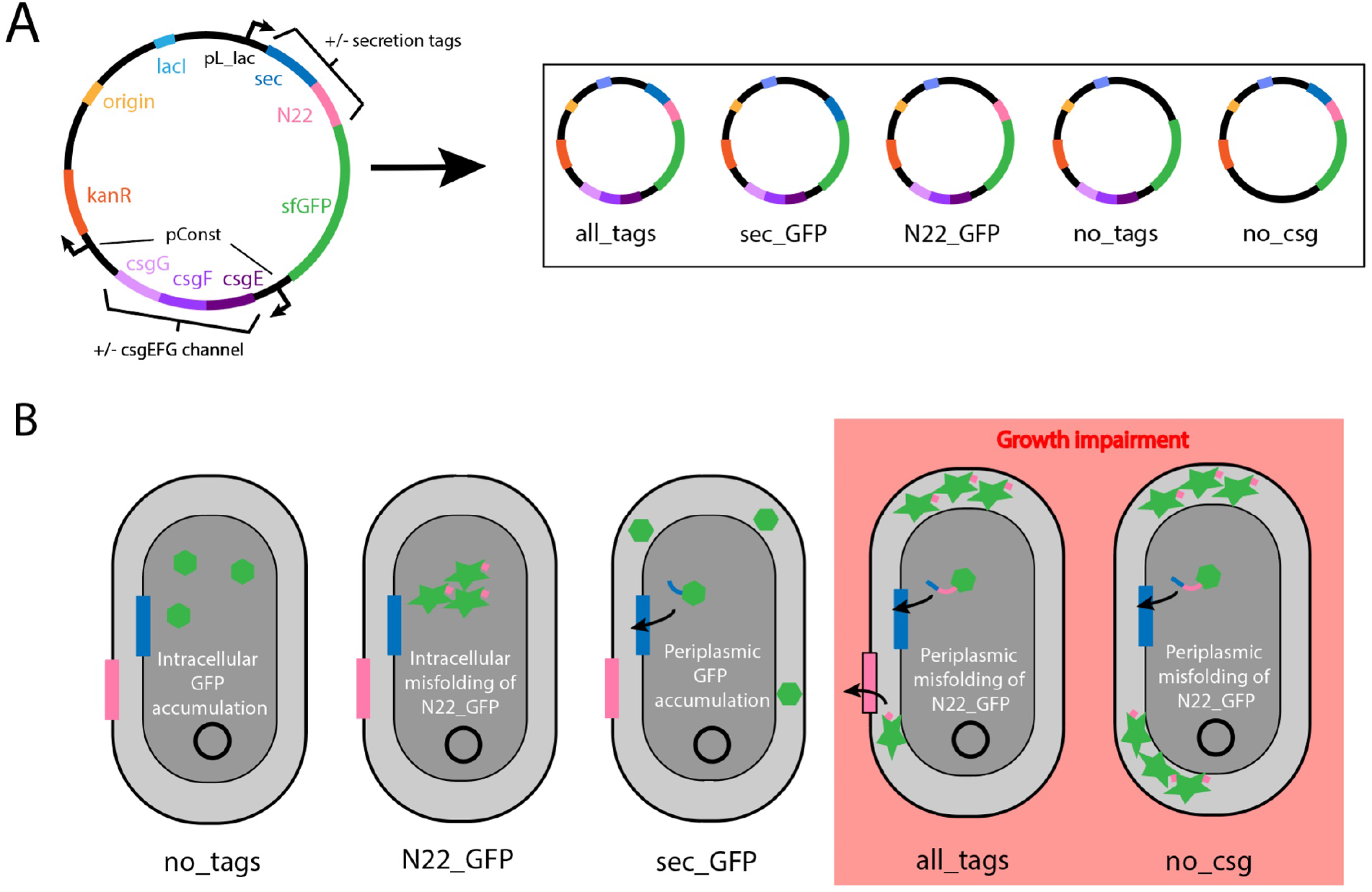
A plasmid was designed with GFP substituted for curli csgA monomer under an inducible promoter and outer membrane channel proteins csgEFG under a constitutive promoter. Permutations with different secretion tags and csgEFG channel protein expression were created (A). Our results suggest that growth impairment is caused by N22_GFP accumulation in the periplasm, leading to protein misfolding and a cellular stress response (B).

In 24-hour kinetic growth experiments, the total GFP fluorescence and absorbance at 600 nm of a culture batch were assayed at various inducer concentrations between 0 and 100 μM, measuring total sfGFP production and cellular density, respectively. The total sfGFP production in batch culture reached its maximum at intermediate levels of induction (Figure 2A), as increasing levels of induction correlated with declining maximum optical density (Figure 2B). Normalizing for cell density, the average growth rate during the first 300 min of growth dropped with increasing IPTG concentration, while average per-cell protein production increased (Figure SI3). The growth-production relationship demonstrates a fundamental tradeoff between these two processes and explains the achievement of a protein production maximum at moderate induction levels (Figure 2C). A propidium iodide stain performed after 24 hours of growth under 10 μM IPTG induction demonstrates significantly higher cell death when GFP is secreted across both membranes compared to intracellular production, suggesting that cellular toxicity from protein secretion is a major source of growth rate impairment, as opposed to nutrient limitation or increased division time (Figure 2D). This is supported by the decreasing slope of the growth curves (Figure 2B) at >400 minutes for higher inducer concentrations.

**Figure 2.**
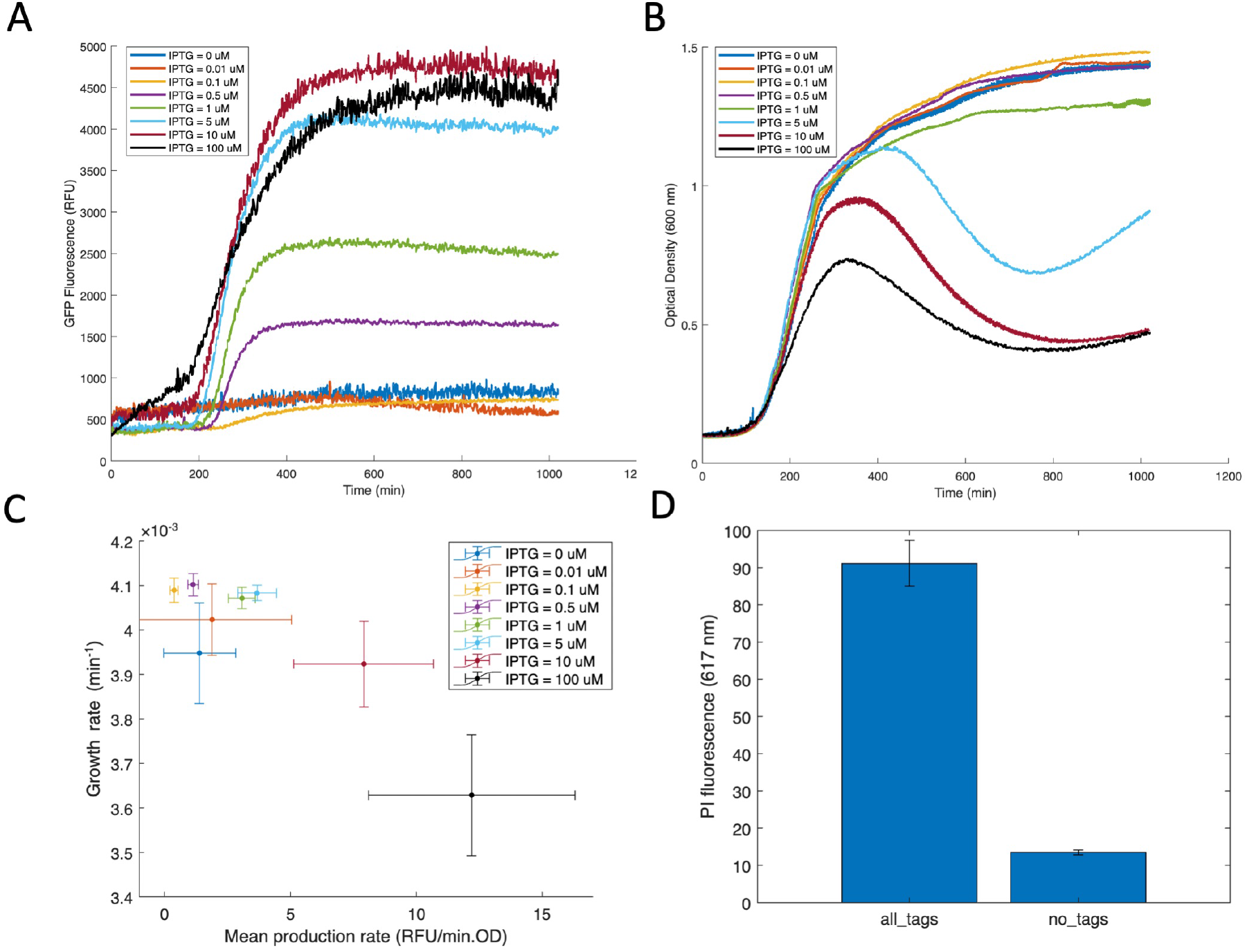
GFP production (A) and cell growth (B) kinetic assays for PBP8 transformed with the all_tags plasmid at various concentrations of inducer. C) Empirical analysis of mean per cell production of GFP versus mean growth rate across all inducer concentrations. D) Propidium iodide staining, showing the relative abundance of dead cells between all_tags and no_tags conditions. Cultures were halted at 24 hours after induction and subjected to the staining protocol. All data expressed as mean values (n=3) with error bars showing standard deviation.

In order to determine the steps within the secretion cascade which contributed significantly to growth impairment, we built and tested a set of plasmid variants that removed one or both of the secretion tags from the N-terminus of sfGFP (Figure 1A), and repeated kinetic growth experiments with these plasmid variants (Figure 3A-B).

**Figure 3.**
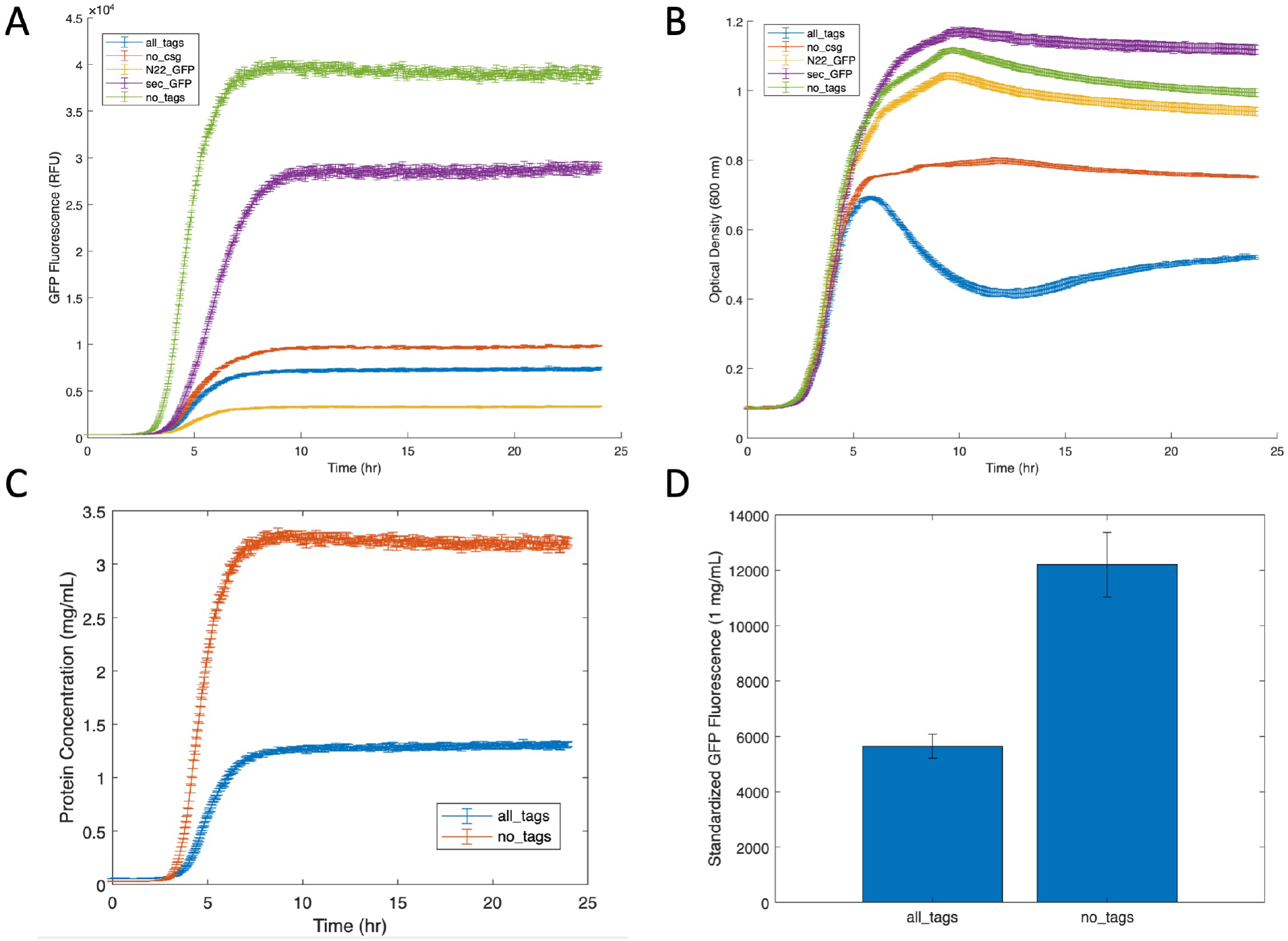
GFP production (A) and cell growth (B) for PBP8 transformed with plasmids encoding GFP with different combinations of N-terminal secretion tags and csg channel proteins (10 μM IPTG). Protein production comparison between all_tags and no_tags expressed in mg/mL, using a Bradford assay conversion from data in Figure 3A. Inset shows standardized RFU/concentration conversion for N22_GFP and no_tags GFP (C). All data expressed as mean values (n=3) with error bars showing standard deviation.

Examining the growth curves of cells harboring each of these plasmids, we noted that the only plasmids causing significant growth depression were those containing both the sec (inner membrane) and the N22 (outer membrane) tags, namely all_tags and no_csg (Figure 3B). Overall, the growth performance of secretion plasmid variants suggests that the observed reduction in cell growth is likely due to periplasmic accumulation of N22_GFP in highly secreting cells. The variant with no secretion tags (no_tags) experienced no growth impairment, indicating that cytosolic accumulation of untagged GFP likely does not impair growth and that high concentrations of IPTG are not inherently cytotoxic (Figure 3B). The unaffected growth of the sec_GFP variant indicates that secretion through the inner membrane and periplasmic accumulation of GFP without N22 do not cause growth impairment (Figure 3B). The unaffected growth of the N22_GFP variant indicates that cytosolic accumulation of N22_GFP does not impair growth (Figure 3C). The only pathological state occurs in the variant when both secretion tags are expressed (all_tags), presumably as N22 GFP accumulates in the periplasm and is secreted via the outer membrane channel.

In order to determine the effect of outer membrane transport versus periplasmic accumulation on growth depression, we characterized PBP8 cells harboring plasmids without csgEFG (no_csg). Compared to the all_tags variant with intact csgEFG, the no_csg variant shows a similar average growth rate during exponential phase and a mild increase in total protein production (Figure 3A-B). Notably, the *csgEFG* deletion was unable to restore growth to levels of the no_tags variant, indicating that the presence of N22_GFP in the periplasm significantly contributes to cell stress and growth depression as opposed to outer membrane transport alone. However, the no_csg variant mildly improved growth compared to all_tags especially in late exponential phase, indicating that outer membrane transport may play a role in reduced cell viability.

When comparing fluorescence results between different plasmid variants (Figure 3A), we found that it was important to account for the differences in fluorescence caused by secretion tag fusion to the GFP protein. Fluorescence measurements were converted to estimated protein concentrations using a Bradford assay standard curve for the all_tags and no_tags GFP variants (Figure 3c). Notably, GFP protein with sec and N22 had lower fluorescence (50-60%) for the same protein concentration compared to untagged GFP, indicating that secretion tags likely disrupt native GFP folding (Figure 3D). While this effect was not measured on N22_GFP (without sec), it provides a possible explanation for the low fluorescence signal of the N22_GFP variant despite the presence of a strong cytosolic band on western blot (Figure 3A, 4B). Even with this adjustment, the all_tags variant demonstrates decreased absolute protein yield compared to no_tags, reaffirming the observation of cellular stress caused by the process of secretion (Figure 3D).

Fractionation studies performed at the end of 24-hour culture experiments allowed for a more nuanced interrogation of the mechanisms leading to periplasmic accumulation (Figure 4A). As expected, the all_tags variant secreted a larger fraction of produced GFP compared to the other variants. It is worth noting that the variants without one or either of the secretion tags secreted a nontrivial amount of produced GFP. These results indicate that the secretion process is likely leaky and nonspecific, complicating the interpretations of sec and N22 as strict inner/outer membrane secretions tags. A western blot assay on the cellular fractions demonstrates the presence of an inclusion body fraction in the induced all_tags variant but not the uninduced or N22_GFP variants, suggesting that inclusion body formation may be a potential reaction to cell stress caused by accumulation of periplasmic protein (Figure 4B).

**Figure 4.**
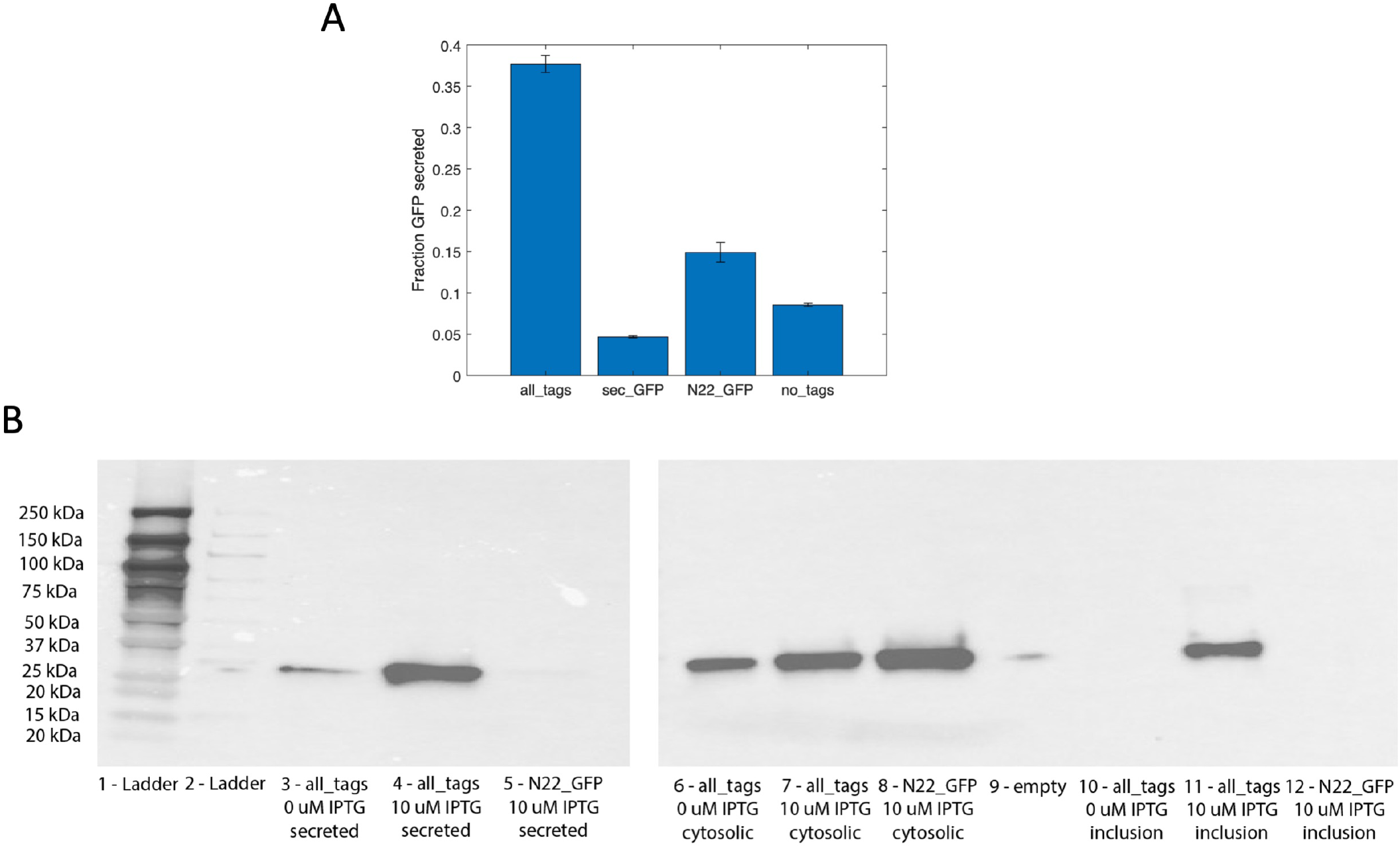
Analysis of GFP compartmentalization in various cellular fractions for GFP bearing different secretion tag combinations. Secreted GFP fluorescence (from the supernatant) expressed as a fraction of total GFP fluorescence for GFP variants (A). Western Blot analysis of GFP concentration in different cellular fractions (secreted, cytosolic, inclusion bodies) and induction conditions (0 or 10 μM IPTG) (B). Data expressed as mean values (n=3) with error bars showing standard deviation.

RNA sequencing data from batch culture experiments comparing the all_tags and N22_GFP variants provides further evidence of periplasmic stress as a mechanism for growth rate depression. A summary of differential gene expression is given in Table 1 and the names and functions of the most highly differentially expressed genes are given in Table 2. Unlike the N22_GFP induced variant, the all_tags induced variant produced a markedly different expression profile from the uninduced control (Table 1, Figure 5A). The top five differentially expressed transcripts in the all_tags variant compared to the N22_GFP variant all correspond to the phage shock protein (PSP) operon, a well-characterized periplasmic stress response gene in *E. coli* (Table 2, Figure 5B). Significantly increased expression of these genes in the all_tags variant compared to N22_GFP indicates that the presence of protein in the periplasm may lead to periplasmic stress and cell death. Gene ontology analysis on the cohort of differentially expressed genes reveals a significant enrichment of genes relating to oxidation-reduction chemistry (Table 1), consistent with known stress response pathways.

**Table 1.**
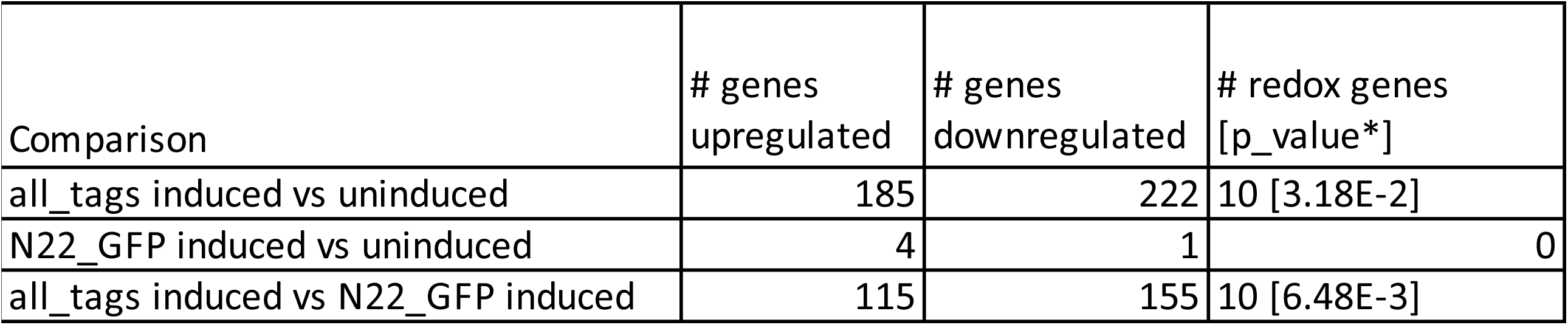
Comparison of the quantity of upregulated and downregulated genes (p_adj < 0.01 and fold change > 1) in the induced and uninduced conditions, with specific subanalysis of redox genes. P-value calculated using Fisher’s exact test from the Panther gene ontology database.

**Table 2.** Top upregulated and downregulated in the all_tags induced variant when compared to N22_GFP. Gene function derived from Uniprot online gene function database [24].

| Gene | Log2 fold change | p-value | Gene Function* |
| --- | --- | --- | --- |
| GFP | 7.97 | 6.89E-23 | sfGFP expressed via the pSecGFP1h plasmids. Contains the Sec and N22 signal sequences used by csgA. |
| pspG | 5.91 | 1.68E-98 | Effector of the phage shock response; upregulated by pIV secretin stress. |
| pspA | 5.73 | 2.83E-120 | Regulatory protein for phage-shock-protein operon. |
| pspB | 5.71 | 1.57E-109 | Psp operon transcription co-activator. |
| pspC | 5.58 | 1.54E-143 | Psp operon transcription co-activator. |
| pspD | 5.47 | 4.54E-137 | Peripheral inner membrane phage-shock protein. |
| rstA | 3.89 | 1.8E-25 | Has roles in the regulation of virulence, acid tolerance, and biofilm formation. |
| mgrB | 3.82 | 1.1E-33 | Represses PhoP/PhoQ signaling, possibly by binding to the periplasmic domain of PhoQ. |
| mgtT | 3.65 | 3.54E-20 | Small protein, function not well understood. |
| norV | 3.33 | 1.04E-21 | NorV and NorW constitute a nitric oxide reductase that couples NADH oxidation to NO reduction. |
| ycjM | 3.32 | 2.21E-18 | May be a regulator of intracellular levels of glucosylglycerate. |
| clbJ | -3.47 | 6.01E-14 | Peptide synthetase in the colibactin biosynthesis pathway. |
| clbG | -3.5 | 8.53E-14 | Acyltransferase in the colibactin biosynthesis pathway. |
| clbL | -3.56 | 4.53E-15 | Amidase in the colibactin biosynthesis pathway. |
| ompC | -3.74 | 4.96E-14 | Forms pores that allow passive diffusion of small molecules across the outer membrane. |
| clbI | -3.78 | 1.67E-14 | Polyketide synthase in the colibactin biosynthesis pathway. |
| clbF | -3.86 | 1.33E-14 | Dehydrogenase in the colibactin biosynthesis pathway. |
| clbE | -3.96 | 1.58E-14 | Phosphopantetheine attachment site in colibactin biosynthesis pathway. |
| clbD | -4.08 | 6.81E-15 | Dehydrogenase in the colibactin biosynthesis pathway. |
| malE | -4.08 | 5.24E-10 | Maltose/maltodextrin transport system substrate-binding protein; part of the ABC transporter complex. |

**Figure 5.**
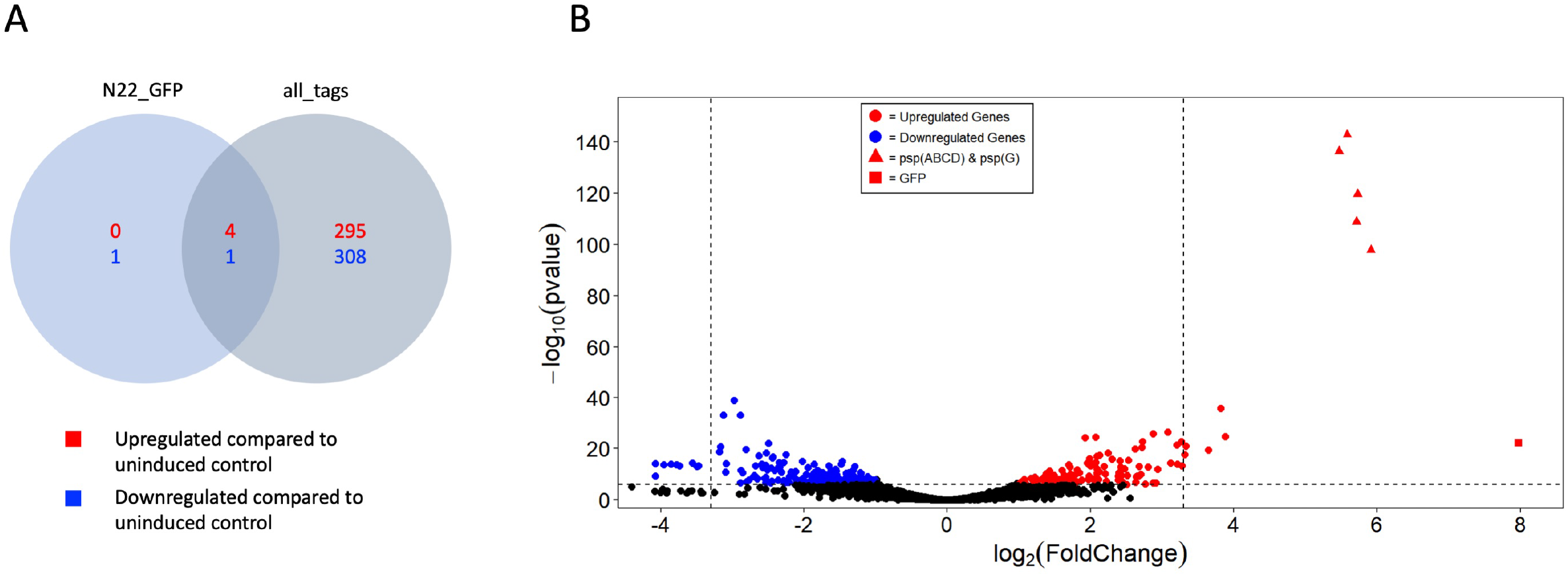
RNA-seq data comparing PBP8 bearing the all_tags plasmid (induced or uninduced) and the N22_GFP plasmid. Comparison of the quantity of upregulated and downregulated genes (p_adj < 0.01 and fold change > 1) in each of the induced conditions, with respect to the uninduced condition (A). Volcano plot showing log10(p-value) as a function of log2-fold change (B). Colors indicate cut-offs for inclusion in 5A. Dotted lines indicate cut-offs for inclusion in Table 2.

## Discussion

In this paper, we empirically describe the relationship between cell growth and secretion of GFP in *E. coli* Nissile 1917 using the Type VIII (curli) secretion apparatus. Our results demonstrate that protein secretion and cell growth are inversely related especially at high inducer concentrations, and this leads to maximum GFP production at intermediate inducer concentrations. Through a series of experiments imposing combinatorial changes to the secretion system, we conclude of that growth depression is likely a stress response to periplasmic accumulation of N22_GFP in cells with a high secretion load.

The deleterious effects of the N22 signal tag on protein function have been previously explored. Since N22 is part of an amyloid protein monomer, it seems to display a tendency to form unstable aggregates when expressed in high quantities [14]. Our results confirm that sfGFP fluorescence is reduced by up to 60% with the N22 tag present, suggesting a disruption of protein folding caused by the N-terminal sequence. Furthermore, the potential instability of N22_GFP suggests a possible mechanism for stress in response to periplasmic protein accumulation. Accumulation of unstable protein aggregates in the periplasm seems to trigger a stress response which leads to the formation of inclusion bodies and contributes to growth impairment. When the inner membrane secretion tag is not expressed, N22 GFP seems to be appropriately sequestered in the cytosol (although less fluorescent), and this stress response is not noted. The difference in redox environments in the cytosol and periplasm, as well as the presence of different protein chaperones may explain the cytosol’s increased ability to sequester these unstable protein products [15, 16].

RNA sequencing data provide further evidence that periplasmic stress may mediate the reduction in cell growth for highly secreting strains. The all_tags variant which secretes GFP (bearing the N22 sequence) into the periplasm demonstrates markedly increased expression of genes in the phage shock protein (PSP) operon compared to a variant which produces cytosolic N22_GFP, as 5 of the 6 most differentially expressed genes are from this single operon. The PSP operon has been characterized as part of a potent stress-response pathway, inducible by heat shock and osmotic shock and leading to more robust stationary phase survival in resource-limited conditions. Moreover, these genes have known inner membrane and periplasmic localization, suggesting a periplasmic stress response to N22_GFP that may mediate growth depression [17].

Overall, our results motivate strategies for engineering stable, high yielding bacterial secretory systems. First, it is clear that the deleterious effects of secretion are linked with the presence of the N22 signal sequence, which means that these same tradeoffs may not be present with secretion using other outer membrane secretion designs. Changing the outer membrane signal sequence may improve protein stability and prevent harmful accumulation in the periplasm. Some studies have previously demonstrated tradeoffs between growth and extracellular protein transport in phylogenetic analyses, but experimentation is needed to better understand the generalizability of the growth-secretion tradeoff [18].

Second, preventing accumulation in the periplasm may be achieved through better regulation of the inflow and outflow of protein into the periplasmic space. N22_GFP accumulation in the periplasm seems to cause growth inhibition, indicating that strategies to reduce the rate of inner membrane transport (e.g., lower sec channel expression, less active inner membrane secretion tag) may lead to increased long-term protein output. As removal of the CsgEFG channel apparatus seemed to further exacerbate growth depression, perhaps increasing the expression or activity of csg could decrease the periplasmic load and improve secretion. Finally, feedback mechanisms which respond to cell stress could further increases the scope of optimization in these secretion systems. Our results from RNA sequencing point to specific pathways that maybe useful targets for mitigating maladaptive stress response for secretion systems.

## Materials and Methods

### Strain and plasmid Engineering

All assays were performed in the *E. coli* Nissle 1917 (EcN) derived PBP8 cells. The PBP8 strain differs from the EcN genome only by the deletion of *csgBAC* and *csgDEFG* operons, which have been replaced by a chloramphenicol resistance gene [19].

Plasmids were constructed using the plasmid backbone from pL6FO encoding sfGFP fused to the N-terminal secretion tags originating from the *csgA* sequence, under lac inducible promotion, *csgEFG* under constitutive promotion, and a kanamycin resistance gene (Figure 1) [20]. The secretion tag permutations and *csgEFG* sequences were originally obtained from plasmid pL6FO and match the *E. coli* MC4100 genomic sequence. Plasmids were constructed by Gibson Assembly, with DNA fragments obtained by PCR with the Q5 polymerase (NEB) [21]. Cloning was performed in the Mach1 *E. coli* strain (Thermo Fisher Scientific) using chemically competent cells, selecting on plates with 50 µg/ml kanamycin. Plasmids were then harvested using Qiagen Miniprep kit and sent for confirmatory Sanger sequencing with GeneWiz.

Each plasmid variant was transformed into PBP8 cells using electroporation. Electrocompetent PBP8 cells were mixed with plasmid DNA before being subjected to a 1250 V pulse. The cells were immediately transferred to SOC growth media and were cultured for 1 hour at 37°C and 225 RPM before being plated on LB agar plates containing 50 µg/ml kanamycin.

### Cell Culture

Cells were cultured in a shaking incubator at 37°C and 225 RPM for the duration of the specified times. All cell cultures contained 50 μg/mL kanamycin to select for cells containing the engineered plasmids. Single colonies taken from LB agar plates were used to make 2 mL uninduced starter cultures that were grown overnight for all experiments. Cells were then diluted 1000-fold into fresh media for culturing in secretion assays.

### Kinetic Assays

A SpectraMax microplate reader (Molecular Devices) was programmed to run kinetics assays to track sfGFP fluorescence and OD_600_ values over time. PBP8 cells from starter cultures were diluted in LB broth in 24-well plates and induced just prior to the 0-hour timepoint at the specified IPTG concentrations. Cells were grown at 37°C with periodic shaking in between fluorescence and OD_600_ measurements, which each occurred every 3 minutes. To track sfGFP fluorescence, the sample excitation and emission wavelengths were set to 488 nm and 510 nm, respectively. To track OD_600_, the absorbance of light at 600 nm was measured. All samples were run in triplicate with mean values being plotted in figures.

### Fractionation

Induced PBP8 cells containing engineered plasmid variants were spun down in a centrifuge at 4000 x g and 4°C for 10 minutes after 24 hours of growth. The supernatant was taken to be the secreted sfGFP fraction. A periplasmic extraction was performed using a 50mM Tris/0.53 M EDTA/20% Sucrose (TES) buffer based on a protocol optimized by Ghamghami et al [22]. Cell pellets were resuspended in the 0.5 mL of cold TES buffer and were incubated on ice for 1 hour. The suspensions were spun down at 15,000 x g at 4°C for 30 minutes and the supernatant was taken to be the periplasmic sfGFP fraction. Cell pellets were resuspended in a 20mM Tris/500mM NaCl/10% Glycerol buffer containing Bugbuster lysis reagent (Millipore Sigma). The suspensions were spun down at 8,000 x g at 4°C for 10 minutes and the supernatant was taken to be the cytosolic sfGFP fraction. The procedure for the isolation of inclusion bodies was adapted from a Biologics International Corporation protocol [23]. The pellet containing inclusion bodies was resuspended in 1 mL 2% Triton X-100 detergent (Sigma-Aldrich) to solubilize membranes. The suspension was spun down at 8,000 x g at 4°C for 20 minutes and the supernatant was discarded as cell debris. The cell pellet was resuspended in 8 M urea to solubilize sfGFP from the inclusion bodies.

### Western Blots

sfGFP isolated from each fraction (secreted, periplasmic, cytoplasmic, and inclusion body) was diluted in a 20mM Tris/500mM NaCl/10% Glycerol buffer such that the volume of each fraction was equivalent to the original culture volume to accurately assess the sfGFP concentration of each fraction in the cultured cells. The fractions were then mixed with Laemmli loading buffer and 100 mM dithiothreitol and heated at 70°C for 10 minutes. Samples were then subjected to SDS-PAGE. Proteins in the gels were transferred to PVDF membranes using the iBlot 2 transfer device (Invitrogen) and corresponding protocol. Membranes were then washed in 1X TBST and placed in a 5% BSA blocking solution for 1 hour. Membranes were then incubated with an anti-His antibody with HRP conjugate (Invitrogen) in 1% BSA at 4°C overnight with gentle rocking. Membranes were then washed 3 times with 1X TBST for 10 minutes each before being transferred to a solution containing enhanced chemiluminescence (ECL) substrate to allow visualization of His-tagged sfGFP (Bio-Rad). The blotted proteins were visualized in a ChemiDoc Imaging System (Bio-Rad) using a chemiluminescence detector setting.

### GFP Purification and Bradford Assay

PBP8 cells containing engineered plasmid variants were cultured in 250 mL LB broth and induced with 10 μM IPTG overnight. Cells were spun down at 4,000 x g at 4°C for 10 minutes before being resuspended in PBS containing 1X Bugbuster lysis reagent (Millipore Sigma). Suspensions were incubated on ice for 30 minutes before being spun down at 15,000 x g at 4°C for 30 minutes. The supernatant was then flowed through Ni-NTA gravity flow columns (ThermoFisher), allowing binding between His-tagged sfGFP and the Ni-NTA resin. The column was washed with 2 resin volumes of 25 mM imidazole in PBS twice before eluting. Elution was performed an increasing concentrations (40-500 mM)of imidazole in PBS. Pure fractions of sfGFP were identified by SDS-PAGE stained with Coomassie Blue.

Buffer exchange of purified sfGFP into PBS was performed by size exclusion (ThermoFisher) to remove high concentrations of imidazole from samples. A set of several BSA standards (0.01-2 mg/mL) were prepared in a 96-well plate in duplicates. Purified sfGFP samples were added to the plate in duplicates. Coomassie Blue dye reagent (Bio-Rad) was added to each well and the absorbance at 595 nm was measured after 5 minutes of incubation. A plot of absorbance vs. BSA concentration was created to generate a standard curve. The concentrations of the sfGFP samples were determined using their corresponding absorbance values and the standard curve. Each sfGFP sample was diluted to 0.4 mg/mL. The fluorescence values of the diluted samples were measured in triplicate using excitation and emission wavelengths of 488 nm and 510 nm, respectively.

### RNA Sequencing

PBP8 cells containing either the all_tags or N22_GFP encoding plasmids were grown in triplicate in a shaking incubator at 37degC for 5.5 hours (end of log growth phase). The cells were pelleted and then frozen at -80degC. The cell pellets were submitted for RNA-seq preparation and analysis (Azenta.Genewiz). mRNA was extracted and purified before being sequenced using the Illumina HiSeq platform. mRNA sequences were then trimmed and mapped to the *E. coli* Nissle 1917 genome (https://www.ncbi.nlm.nih.gov/nuccore/CP022686.1). Gene hit counts to the reference genome were used for differential gene expression analysis to compare gene expression between the tested groups. The Wald test was used to generate p-values and log2 fold changes for each comparison. P-values were adjusted using a Bonferroni correction corresponding to the number of genes tested. Gene ontology analysis was done using the Panther online database, comparing the differentially expressed genes against the GO molecular function database to identify statistically significant enrichment.

### Data Processing

All data and figures were analyzed in Matlab. For kinetic experiments, average growth rates were calculated using the average slope of a log-linear fit over the exponential growth phase, determined to be around 200 minutes. Protein production rate per cell was calculated by taking the rate of change of fluorescence and dividing by the optical density at a given point in time. The average per cell production rate was calculated over the protein production interval, determined to be around 300 min. Conversions between fluorescence and protein concentration were calculated using the standard curve derived from the Bradford assay.

## Material Availability Statement

All materials including plasmids, bacterial strains, laboratory protocols, and reagent information are available as per the Biobricks Open Material Transfer Agreement (https://biobricks.org/open-material-transfer-agreement/).

## Data Availability Statement

All annotated plasmid sequences are uploaded as GenBank files.

RNA seq data was uploaded to GEO under the ENCODE guidelines (accession #: GSE221524; reviewer token #: ghkbqswuvjubzmn).

All other primary data are submitted as supplemental media.

## Figure Legend

SI1

Kinetic protein production (A) and growth (B) curves at different inducer concentrations for plasmid variant with all secretion tags (all_tags) and csgEFG channel proteins. Data expressed as mean (n=3) with error bars representing standard deviation.

SI2

Kinetic protein production (A) and growth (B) curves at different inducer concentrations for plasmid variant with all secretion tags without csgEFG channel proteins (no_csg).

SI3

Mean growth rate during exponential growth phase decreases with increasing concentration of inducer (A), while mean protein production rate per cell increases with inducer concentration (B). These combined effects lead to a protein production maximum at intermediate inducer concentrations.

