## Supplemental Materials for "Periplasmic stress contributes to a tradeoff between protein secretion and cell growth in E. Coli Nissile"

1a

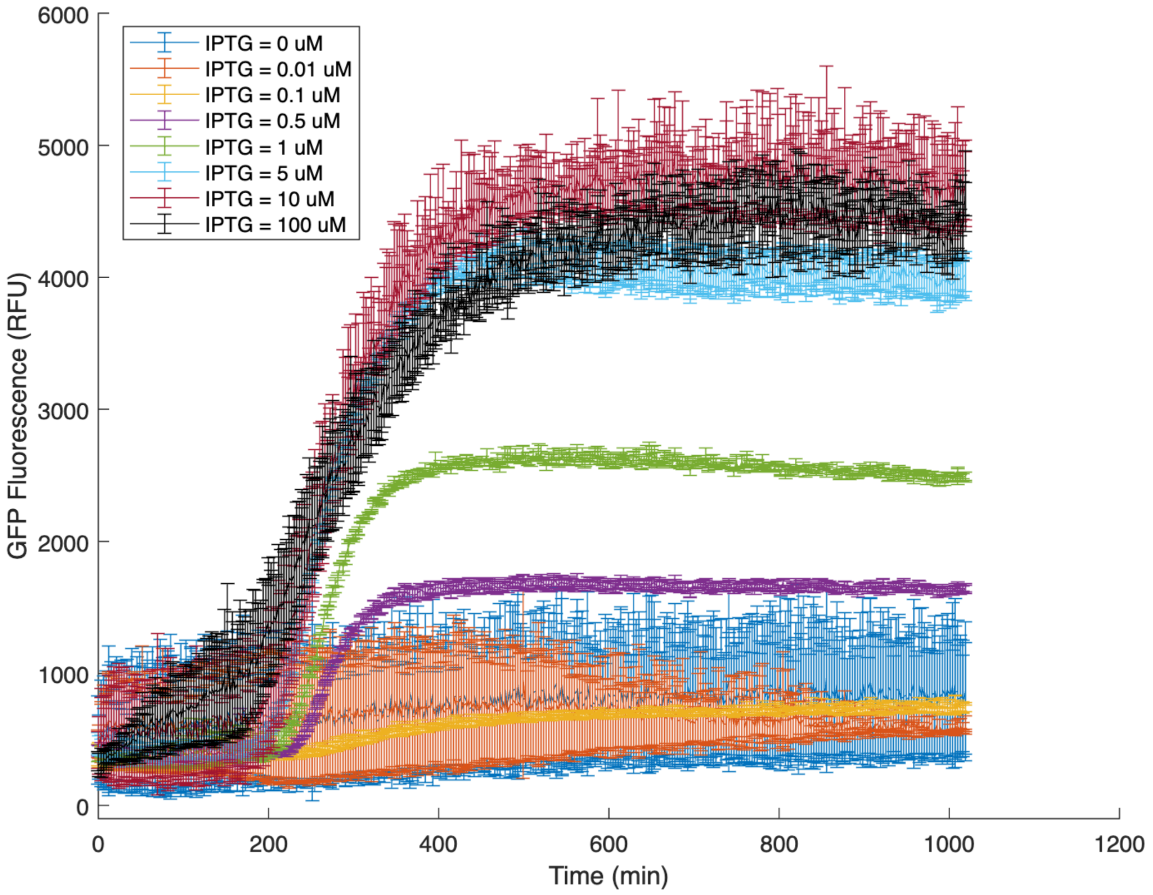

6000
5000
4000
3000
2000
1000
0
GFP Fluorescence (RFU)
0
200
400
600
800
1000
1200
Time (min)
IPTG = 0 uM
IPTG = 0.01 uM
IPTG = 0.1 uM
IPTG = 0.5 uM
IPTG = 1 uM
IPTG = 5 uM
IPTG = 10 uM
IPTG = 100 uM


1b

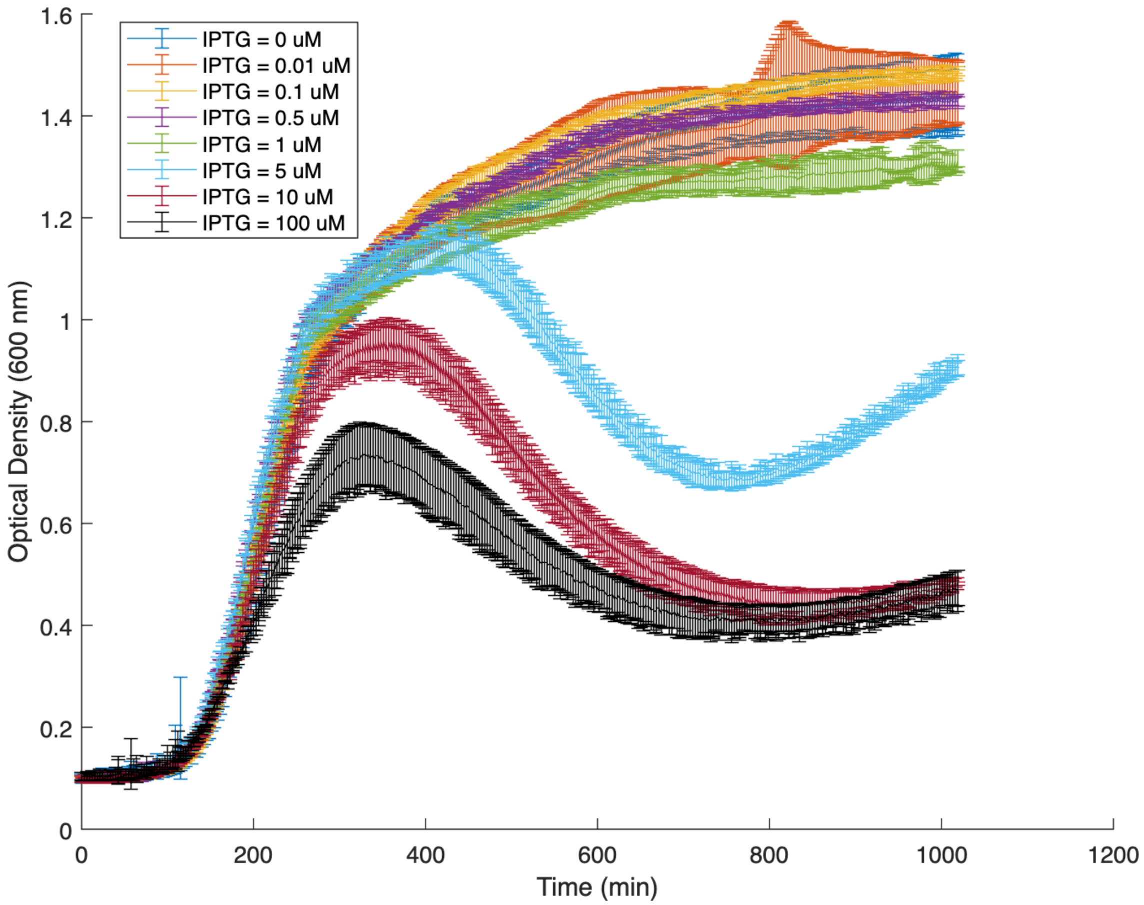

1.6
1.4
1.2
1
0.8
0.6
0.4
0.2
0
Optical Density (600 nm)
0
200
400
600
800
1000
1200
Time (min)
IPTG = 0 uM
IPTG = 0.01 uM
IPTG = 0.1 uM
IPTG = 0.5 uM
IPTG = 1 uM
IPTG = 5 uM
IPTG = 10 uM
IPTG = 100 uM

2a

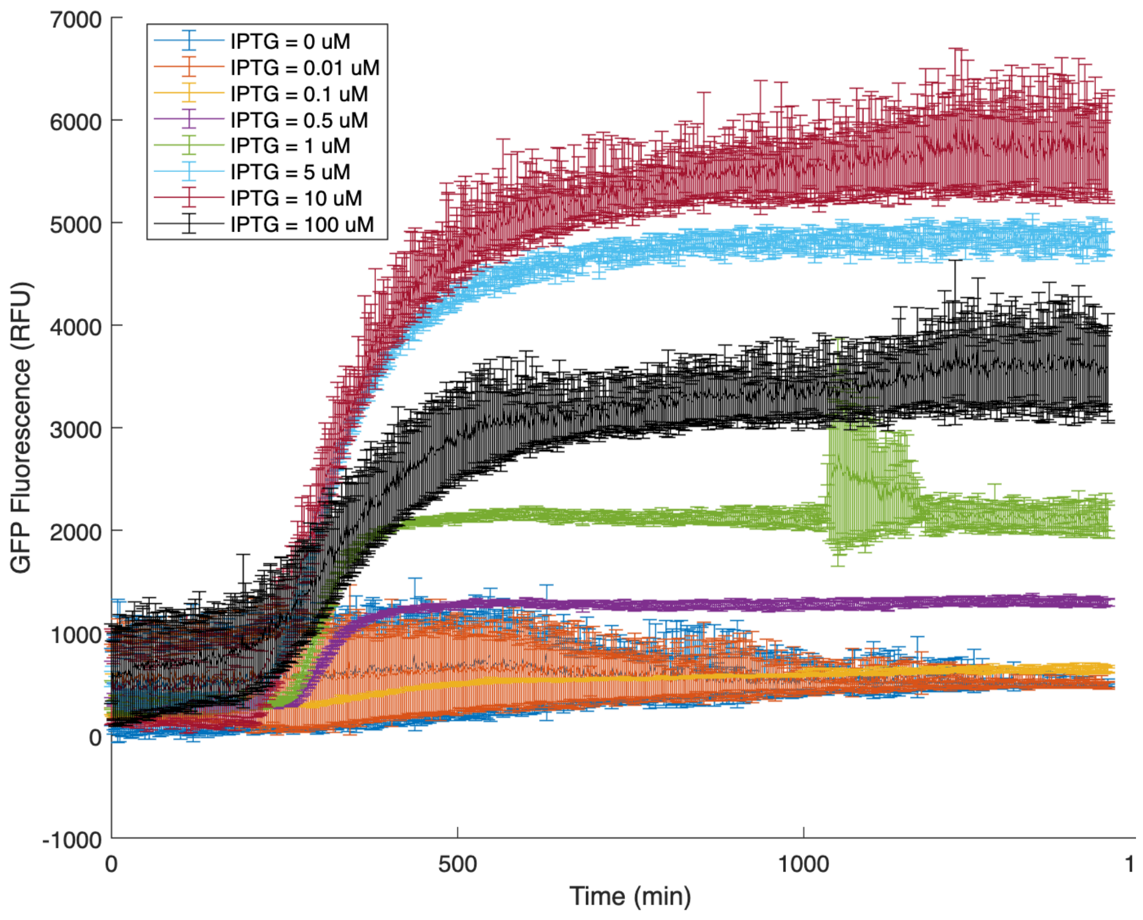
IPTG = 0 uM
IPTG = 0.01 uM
IPTG = 0.1 uM
IPTG = 0.5 uM
IPTG = 1 uM
IPTG = 5 uM
IPTG = 10 uM
IPTG = 100 uM
GFP Fluorescence (RFU)
Time (min)
7000
6000
5000
4000
3000
2000
1000
0
-1000
0
500
1000
1

2b

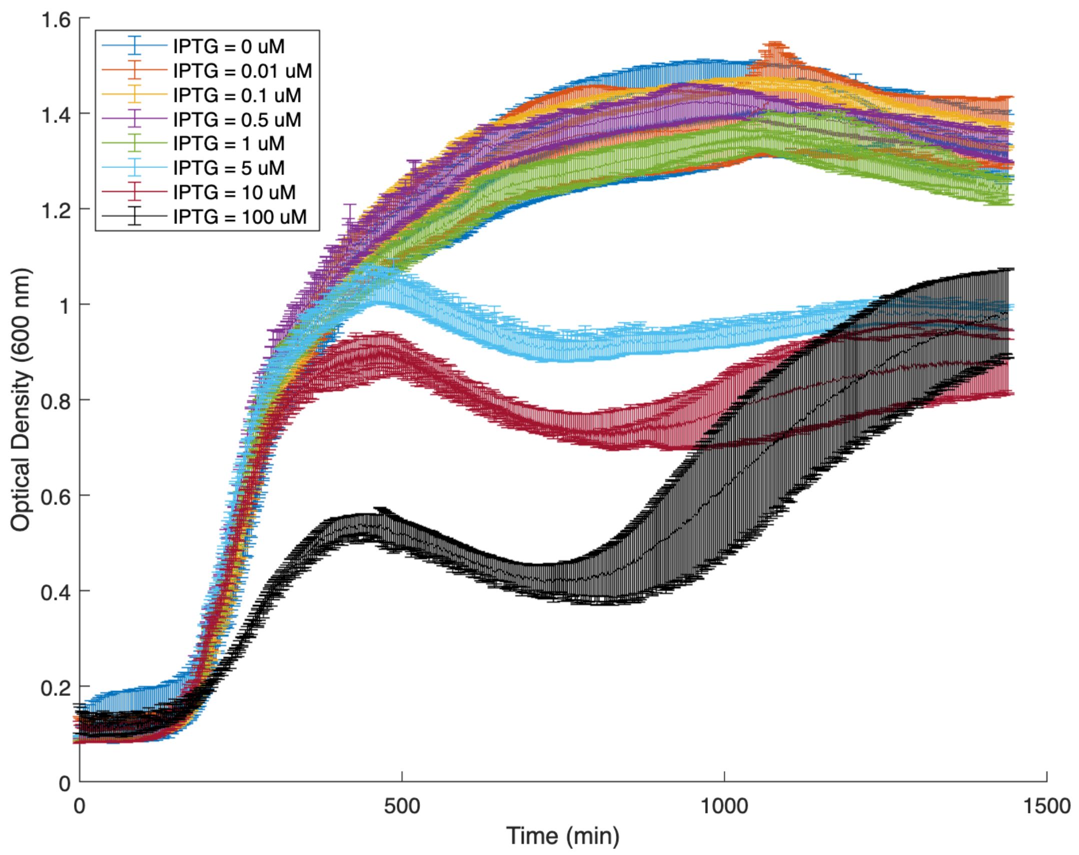
IPTG = 0 uM
IPTG = 0.01 uM
IPTG = 0.1 uM
IPTG = 0.5 uM
IPTG = 1 uM
IPTG = 5 uM
IPTG = 10 uM
IPTG = 100 uM
Optical Density (600 nm)
Time (min)
1.6
1.4
1.2
1
0.8
0.6
0.4
0.2
0
0
500
1000
1500

3a

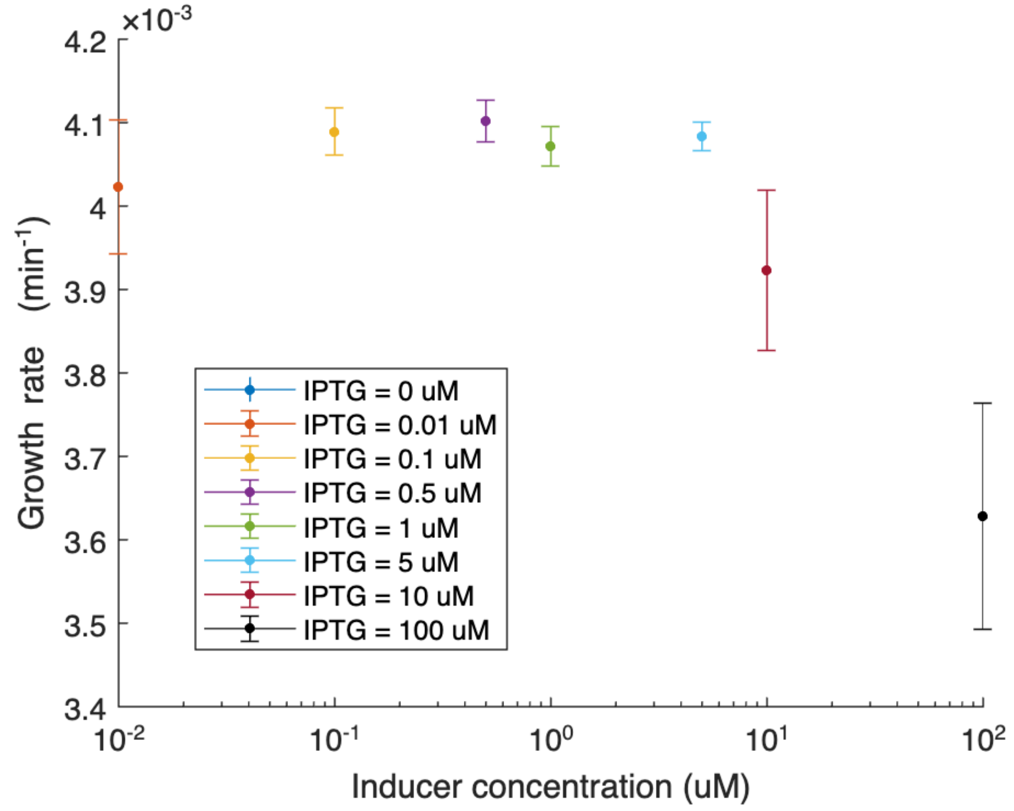
×10-3
Growth rate (min-1)
Inducer concentration (uM)
IPTG = 0 uM
IPTG = 0.01 uM
IPTG = 0.1 uM
IPTG = 0.5 uM
IPTG = 1 uM
IPTG = 5 uM
IPTG = 10 uM
IPTG = 100 uM

3b

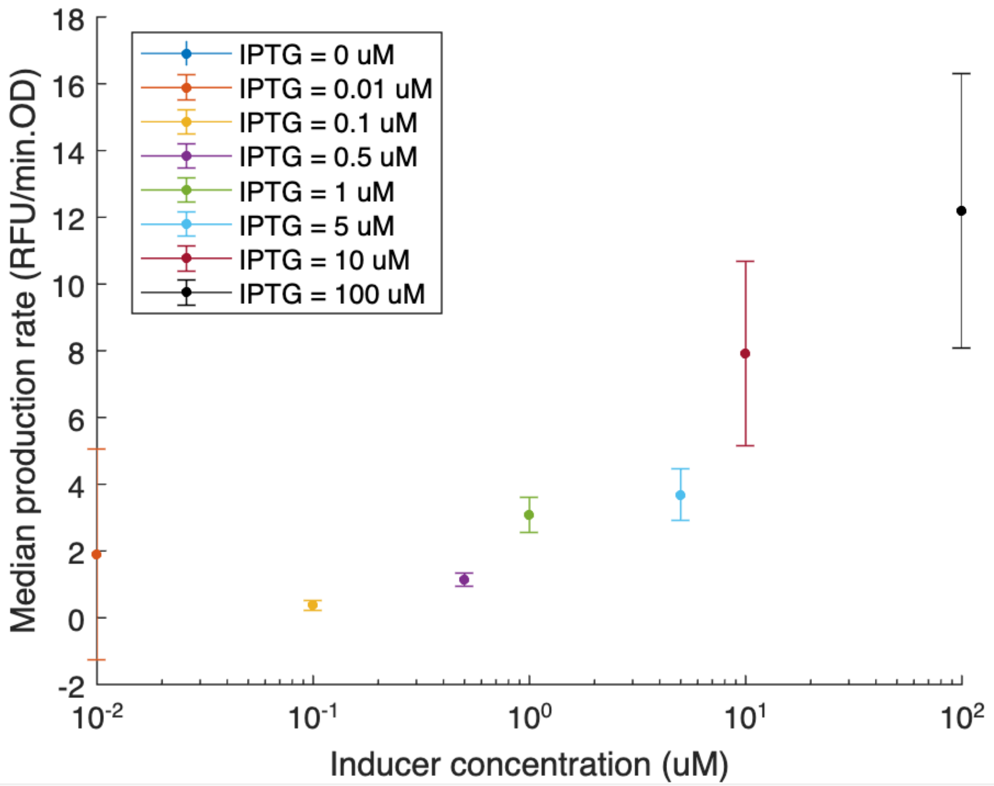
Median production rate (RFU/min.OD)
Inducer concentration (uM)
IPTG = 0 uM
IPTG = 0.01 uM
IPTG = 0.1 uM
IPTG = 0.5 uM
IPTG = 1 uM
IPTG = 5 uM
IPTG = 10 uM
IPTG = 100 uM

### Figure Legends

SI1
Kinetic protein production (A) and growth (B) curves at different inducer concentrations for plasmid variant with all secretion tags (all_tags) and csgEFG channel proteins. Data expressed as mean (n=3) with error bars representing standard deviation.

SI2
Kinetic protein production (A) and growth (B) curves at different inducer concentrations for plasmid variant with all secretion tags without csgEFG channel proteins (no_csg).

SI3
Mean growth rate during exponential growth phase decreases with increasing concentration of inducer (A), while mean protein production rate per cell increases with inducer concentration (B). These combined effects lead to a protein production maximum at intermediate inducer concentrations.
